# The Antiviral and Cancer Genomic DNA Deaminase APOBEC3H Is Regulated by a RNA-Mediated Dimerization Mechanism

**DOI:** 10.1101/203000

**Authors:** Nadine M. Shaban, Ke Shi, Kate V. Lauer, Michael A. Carpenter, Christopher M. Richards, Michael W. Lopresti, Daniel Salamango, Jiayi Wang, Surajit Banerjee, William L. Brown, Hideki Aihara, Reuben S. Harris

## Abstract

Human APOBEC3H and homologous single-stranded DNA cytosine deaminases are unique to mammals. These DNA editing enzymes function in innate immunity by restricting the replication of viruses and transposons. Misregulated APOBEC3H also contributes to cancer mutagenesis. Here we address the role of RNA in APOBEC3H regulation. APOBEC3H co-purifies with RNA as an inactive protein, and RNase A treatment yields enzyme preparations with stronger DNA deaminase activity. RNA binding-defective mutants are DNA hypermutators. Chromatography profiles and crystallographic data demonstrate a mechanism in which double-stranded RNA mediates enzyme dimerization. RNA binding is required for APOBEC3H cytoplasmic localization and for packaging into HIV-1 particles and antiviral activity. Related DNA deaminases including other APOBEC3 family members and the antibody gene diversification enzyme AID also bind RNA and are predicted to have a similar RNA binding motif suggesting mechanistic conservation and relevance to innate and adaptive immunity and to multiple diseases.

**HIGHLIGHTS:** RNA inhibits human APOBEC3H DNA cytosine deaminase activity

RNA binding mutants are DNA hypermutators

X-ray structure demonstrates an RNA duplex-mediated APOBEC3H dimerization mechanism

RNA binding is required for packaging into HIV-1 particles and antiviral activity

## INTRODUCTION

Human cells have the potential to express up to nine active DNA cytosine deaminase enzymes collectively called APOBECs (Conticello, 2008; Harris and Dudley, 2015). Activation-induced deaminase (AID) is expressed in antigen-responsive B lymphocytes, and it initiates the diversification of antibody genes through the distinct processes of somatic hypermutation and class switch recombination (Di Noia and Neuberger, 2007; Robbiani and Nussenzweig, 2013). APOBEC1 is expressed in gastrointestinal tissues, and it edits a range of RNA species including the *APOB* mRNA (motivating the original name for the family) (Fossat and Tam, 2014; Koito and Ikeda, 2013). APOBEC3A (A3A), APOBEC3B (A3B), APOBEC3C (A3C), APOBEC3D (A3D), APOBEC3F (A3F), APOBEC3G (A3G), and APOBEC3H (A3H) are expressed at varying levels in different human tissues, and these enzymes function broadly and inducibly in innate immunity by restricting the replication of DNA-based parasites including common retrotransposons L1 and Alu and retroviruses such as HTLV-1, HIV-1, and HIV-2 (Harris and Dudley, 2015; Malim and Bieniasz, 2012; Simon et al., 2015).

Although single-stranded DNA cytosine deamination is the hallmark activity of the APOBEC enzyme family, RNA is important for several aspects of APOBEC biology. First, as alluded above, RNA editing is a *bona fide* function for at least two family members, APOBEC1 (Teng et al., 1993) and A3A (Sharma et al., 2015). Second, RNA has been implicated in governing the cytoplasmic localization of A3G and A3F, including accumulation in stress granules and RNA processing bodies (Gallois-Montbrun et al., 2007; Izumi et al., 2013; Kozak et al., 2006; Phalora et al., 2012; Stenglein et al., 2008; Wichroski et al., 2006). Third, an RNA-dependent interaction is required for A3G and A3F to interact with HIV-1 Gag, gain access to assembling viral particles, and exert antiviral activity (Apolonia et al., 2015; Bogerd and Cullen, 2008; Cen et al., 2004; Huthoff and Malim, 2007; Khan et al., 2005; Schafer et al., 2004; Svarovskaia et al., 2004; Wang et al., 2007; Wang et al., 2008; York et al., 2016; Zennou et al., 2004). A similar RNA-dependent packaging mechanism is proposed for the other HIV-1 restrictive APOBEC3 family members, A3D and A3H (Wang et al., 2011; York et al., 2016; Zhen et al., 2012). Fourth, the interactions between many different cellular proteins and A3F, A3G, APOBEC1, and AID are disrupted by RNase treatment and therefore likely mediated by RNA (Basu et al., 2011; Chiu et al., 2006; Gallois-Montbrun et al., 2008; Gallois-Montbrun et al., 2007; Kozak et al., 2006; Nowak et al., 2011). Finally, nearly all biochemical studies include RNase treatment as an essential step in APOBEC purification protocols suggesting that RNA may be a negative regulatory factor that competitively prevents these enzymes from accessing single-stranded DNA substrates [*e.g.*, (Ara et al., 2014; Bransteitter et al., 2003; Chelico et al., 2006; McDougall and Smith, 2011; Mitra et al., 2015; Shlyakhtenko et al., 2011; Starrett et al., 2016)]. Despite the overarching importance of RNA in APOBEC biology, mechanistic details are lacking largely due to the absence of molecular and structural information on APOBEC-RNA complexes.

A3H is unique among APOBEC3 subfamily members for several reasons. First, the gene encoding A3H is distinct phylogenetically, shows high conservation, and invariably anchors the 3’-end of the *APOBEC*3 locus (LaRue et al., 2008; Münk et al., 2008). The encoded enzyme exists as a single zinc-coordinating domain Z3-type deaminase in higher primates including humans, or as a single- or double-domain enzyme, typically Z2-Z3 organization, in most other mammals including artiodactyls, carnivores, and rodents (LaRue et al., 2008; Münk et al., 2008). Second, unlike Z1 and Z2-type deaminase genes, the A3H or Z3-type genes show no copy number variation in mammals, existing in each species as a full gene or as the 3’ half of a gene encoding a double domain deaminase. Third, despite strict copy number conservation, A3H is the most polymorphic family member in humans with 7 reported haplotypes due to variations at 5 amino acid positions and 4 reported splice variants (Harari et al., 2009; OhAinle et al., 2008; Wang et al., 2011). These naturally occurring A3H variations are known to influence protein stability, antiviral activity, and subcellular localization, with at least two stable variants capable of potently suppressing Vif-deficient HIV-1 replication (Harari et al., 2009; Hultquist et al., 2011; Nakano et al., 2017; OhAinle et al., 2008; Ooms et al., 2013; Refsland et al., 2014; Wang et al., 2011). A3H is also implicated in restricting the replication of a range of other viruses and transposons (Bouzidi et al., 2016; Hultquist et al., 2011; Kock and Blum, 2008; Ooms et al., 2012; Tan et al., 2009). Moreover, mislocalized A3H (haplotype I) is a likely contributor to cancer mutagenesis (Starrett et al., 2016).

Here we investigate the role of RNA in regulating the activities of A3H. The most common, stable, active, and antiviral A3H variant in humans is the 183 amino acid haplotype II enzyme, hereafter A3H. RNA invariably co-purifies with A3H and suppresses enzyme activity. RNase A treatment removes most, but not all, of the bound RNA and enables strong DNA deaminase activity. Mutagenesis of a predicted positively charged RNA binding patch results in hypermutator activity. Wild-type A3H and hyperactive variants show dimeric and monomeric size exclusion profiles, respectively, suggesting an RNA-mediated dimerization mechanism. Indeed, a crystal structure of a near wild-type human A3H-mCherry protein revealed a duplex RNA-bridged enzyme dimer involving the same amino acid residues that yield DNA deaminase hyperactivity upon mutagenesis. This RNA binding surface is required for A3H cytoplasmic localization and for packaging into HIV-1 particles and antiviral activity. Thus, RNA serves multiple beneficial regulatory roles in A3H biology.

## RESULTS

### RNA Digestion Enhances APOBEC3H Purification and DNA Cytosine Deaminase Activity

Our initial attempts to express and purify His6-SUMO-A3H from *E. coli* were unsuccessful because most protein failed to bind the affinity resin. We speculated that RNA may be promoting aggregation and preventing the hexahistidine tag from binding. We therefore treated lysates with RNase, repeated metal affinity purifications, and compared fractions by SDS PAGE. This analysis revealed a strong recovery of A3H but only with RNase A treatment (**Figure 1A**). Furthermore, using an oligo-based single-stranded DNA deamination activity assay, A3H catalytic activity could only be detected in *E. coli* lysates treated with RNase A suggesting that RNA inhibits deaminase activity (**Figure 1B**). A similar RNase A requirement was found for activity assays of A3H expressed in human 293T cells (**Figure 1C**). Requirements for RNase treatment have been described previously for A3H, as well as for all other family members including A3G and AID, suggesting conservation of the underlying molecular mechanism [*e.g.*, (Bransteitter et al., 2003; Chiu et al., 2005; Gu et al., 2016; Mitra et al., 2015; Shlyakhtenko et al., 2011)].

**Figure 1.**
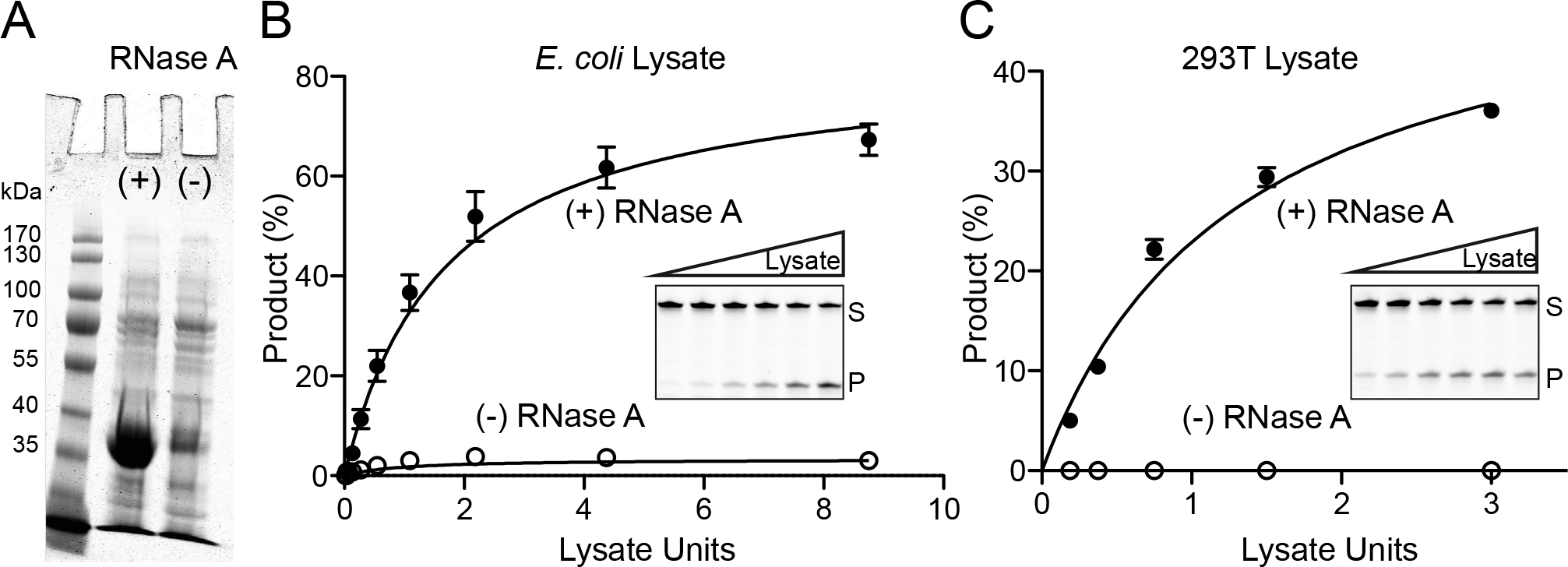
RNase Digestion Enables APOBEC3H Purification and DNA Deaminase Activity. (**A**) Coomassie-stained image showing His6-SUMO-A3H recovery from *E. coli* in the presence and absence of RNase A treatment. (**B**) DNA deaminase activity of His6-SUMO-A3H in extracts from *E. coli* with and without RNase A treatment (mean +/− SD; n = 3 experiments; inset gel image shows A3H-mediated conversion of a single-stranded DNA substrate to product, S to P, in the presence of RNase). Lysate units were chosen to include reactions with single-hit kinetics. (**C**) DNA deaminase activity of untagged A3H in extracts from 293T cells, with experimental parameters similar to those in panel B.

### Identification of Hyperactive APOBEC3H Variants by Altering a Predicted Positively Charged RNA Binding Surface

Since RNA inhibits A3H catalytic activity, we reasoned that RNA may bind the active site pocket and directly prevent the binding of single-stranded DNA. Alternatively, the RNA binding domain may be distinct and indirectly inhibit DNA deamination activity by forming inhibitory complexes. To help distinguish between these possibilities and inform mutagenesis experiments, an A3H structural model was generated (**Figures 2A** and **2B**). Based on recent A3A/B-ssDNA co-crystal structures (Kouno et al., 2017; Shi et al., 2017), A3H loop 1, loop 7, and active site residues define a positively charged region that is likely to be essential for binding single-stranded DNA and may also be involved in binding RNA (*i.e.*, in this model, the two activities will be inseparable by mutagenesis; patch 1 in **Figure 2A**).

**Figure 2.**
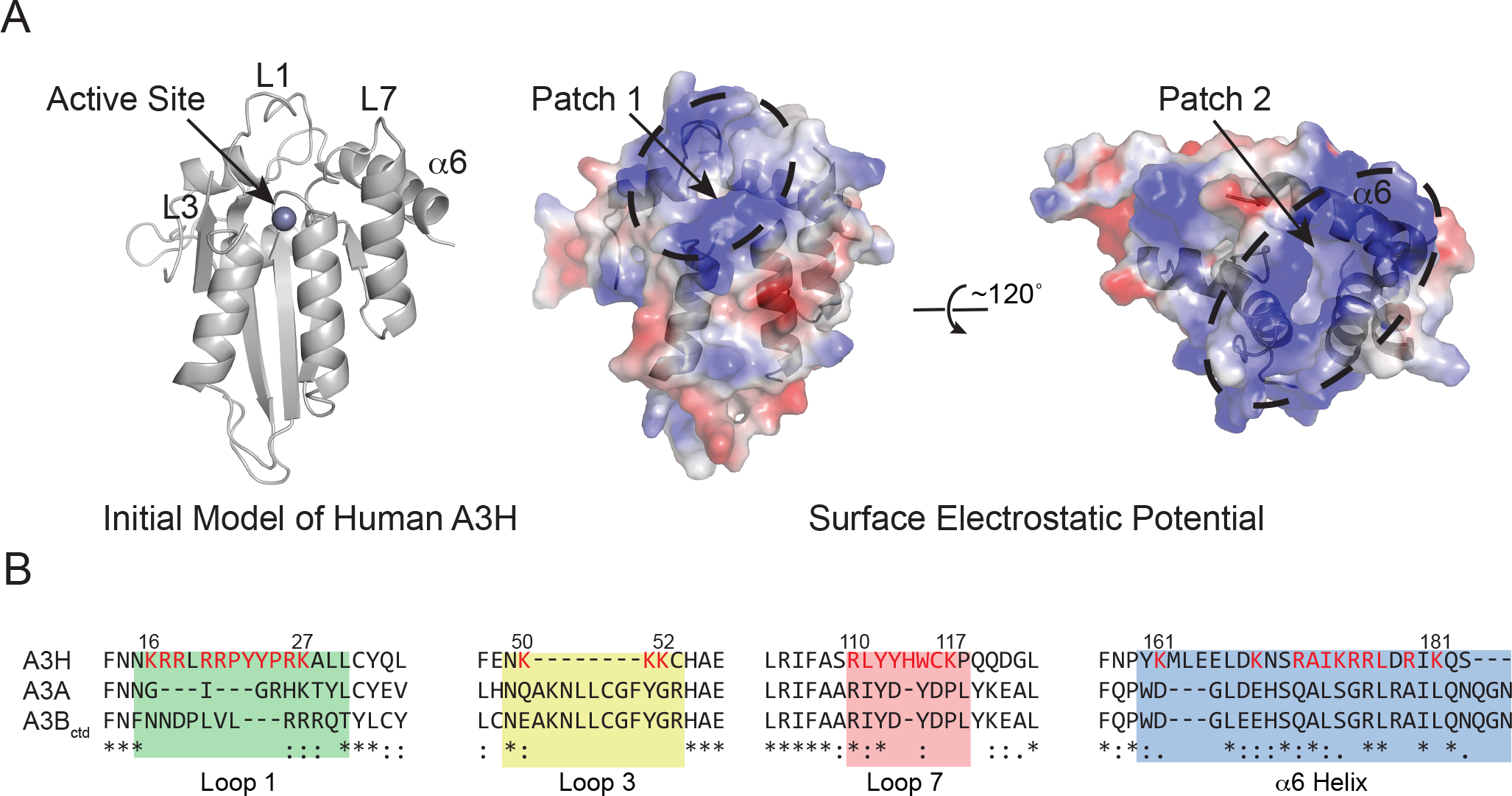
Initial Structural Model Predicts Two Positively Charged Patches in Human APOBEC3H. (**A**) Initial structural model of human A3H generated using Phyre2. The left ribbon schematic shows a single zinc ion in the active site, loops 1, 3, and 7, and the α6-helix. The center and right space-filled schematics show the predicted electrostatic potential for A3H in the same orientation and rotated 120 degrees, respectively. Patch 1 encompasses the active site and residues known to be required for DNA deamination. Patch 2 is distinct and includes several residues in the α6-helix. (**B**) Clustal Omega alignment of human A3H, A3A, and A3B C-terminal domain (ctd) depicting loop 1, loop 3, loop 7, and α6-helix regions. Numbers correspond to the 183 amino acid splice variant of human A3H haplotype II, and residues highlighted in red were changed by site-directed mutagenesis.

Interestingly, the structural model also predicted a second positively charged region centered upon α-helix 6, hereafter referred to as patch 2 (**Figures 2A** and **2B**). Amino acids within this basic patch include Arg171, Lys174, Arg175, Arg176, and Arg179 (Figure 2B). We reasoned that if a residue located in this basic patch is important for binding RNA, then substitutions at this site may impair the formation of RNA-inhibited complexes and simultaneously increase catalytic activity.

A panel of mutant A3H constructs covering both patch 1 and patch 2 was tested in the *E. coli*-based rifampicin-resistance (Rif^R^) mutation assay, which provides a quantitative measure of APOBEC-mediated DNA mutator activity (Harris et al., 2002). Positively charged Arg and Lys residues were changed to Glu, other residues (Leu, Ile, Tyr, Trp) were changed to Ala, and Ala residues were changed to Glu. The background mutation frequency in this assay is approximately 1 × 10^−8^ (empty vector and catalytic mutant E56A negative controls; black dotted line in **Figures 3A** and **3B**). Expression of wild-type A3H caused a 10-fold increase in the median Rif^R^ mutation frequency (red dotted line in **Figures 3A** and **3B**). Several A3H amino acid substitution mutants had no significant affect, such as several loop 1 and 3 changes (R26E, K26E, K50E, K51E, K52E), whereas other substitutions partly or completely abrogated mutator activity (catalytic E56A, loop 1 R21E, P22A, Y23A, Y24A, P25A, and loop 7 R110E, L111A, Y112A, Y113A). The latter group of mutants is most likely defective in DNA deamination (compare with results in **Figure 4**, below).

**Figure 3.**
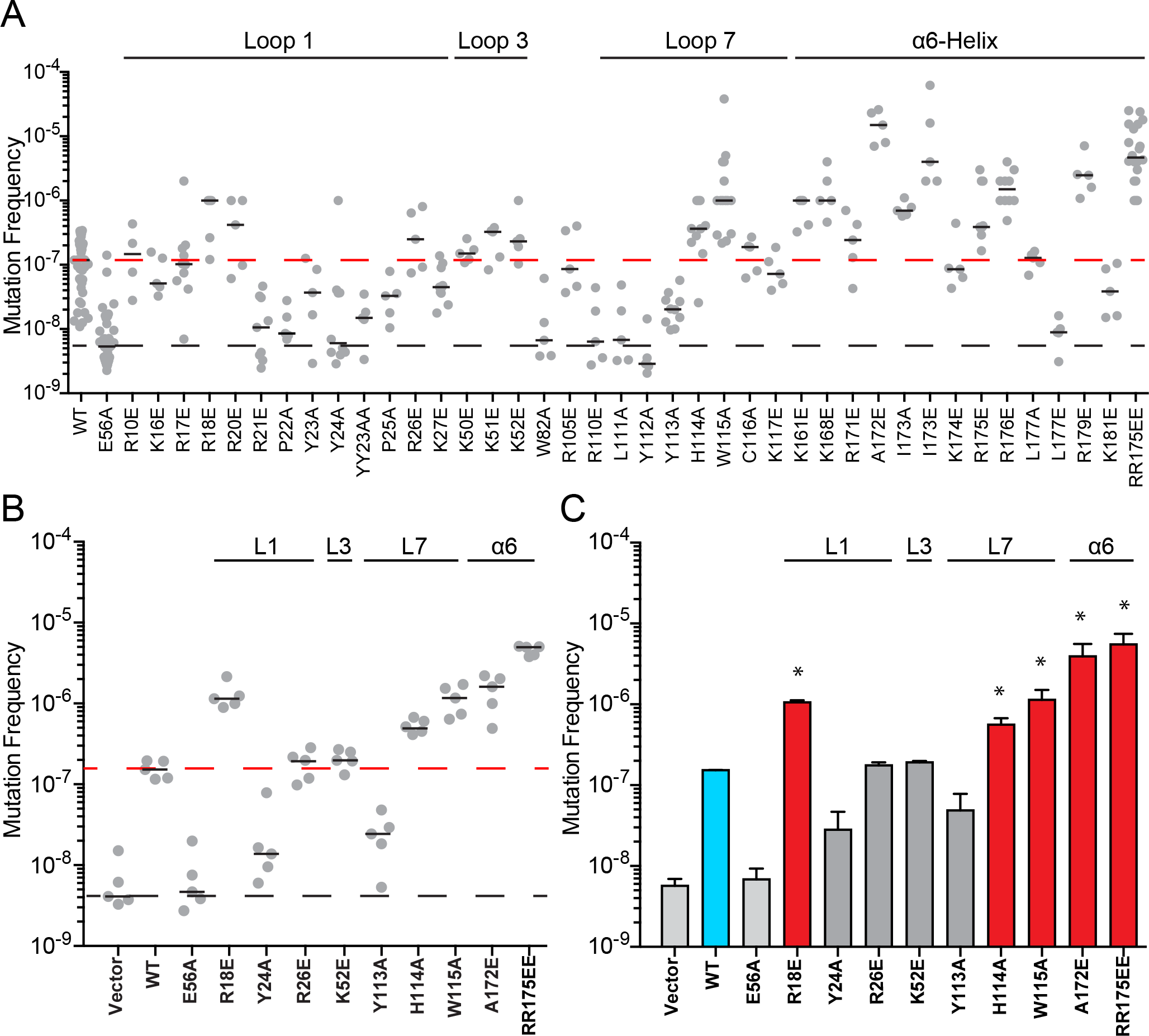
Identification of Hyperactive Human APOBEC3H Mutants. (**A**) Rif^R^ (*rpoB*) mutation frequency of *E. coli* expressing the indicated His6-Sumo-A3H constructs (WT, wild-type, or amino acid substitution mutants). Each dot represents data from an individual culture (n ≥ 5 per condition) with short horizontal lines depicting medians. The black dashed line represents the background mutation frequency of *E. coli* expressing an empty vector (not shown) or A3H catalytic mutant (E56A). The red dashed line shows the mutation frequency of *E. coli* expressing WT A3H. (**B**) Data from a representative Rif^R^ mutation experiment comparing WT A3H and the indicated mutants including 5 hypermutators. Conditions and labels are similar to panel A. (**C**) Quantification of key Rif^R^ mutation data. Each histogram bar reports the mean +/− SEM of the median mutation frequency from 3 independent experiments (panel B and 2 independent experiments not shown). The mutation frequencies induced by the 5 hypermutators (red bars) are significantly greater than that of WT A3H (blue bar; p<0.01 by Student’s t-test).

**Figure 4.**
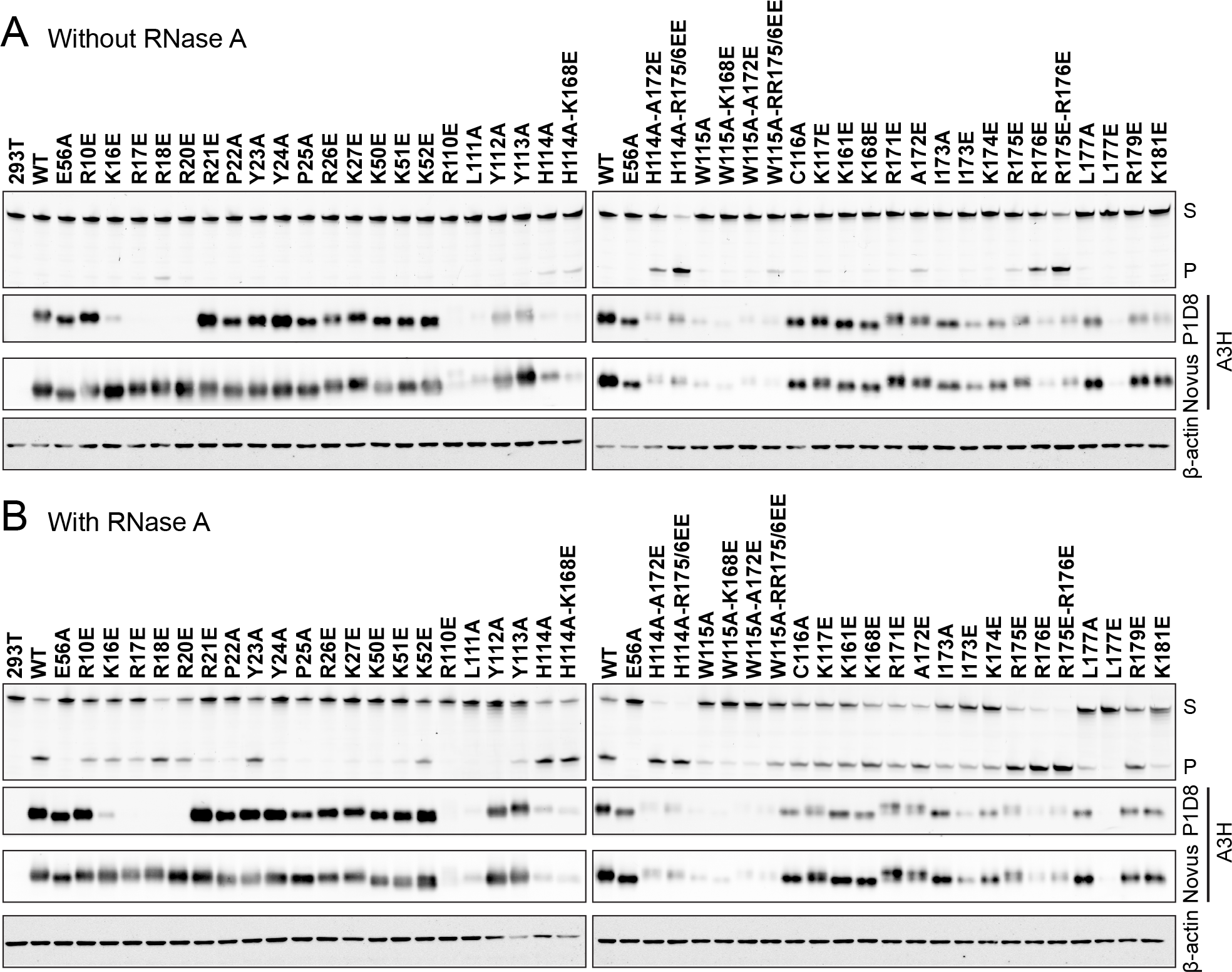
DNA Deaminase Activity of APOBEC3H and Derivatives in 293T Extracts With and Without RNase A Treatment. (**A** and **B**) Comparison of the DNA deaminase activities of untagged wild-type (WT) human A3H or the indicated amino acid substitution mutants in 293T extracts in the absence and presence of RNase (S, substrate; P, product). Corresponding immunoblots show levels of A3H in cell lysates using a murine mAb (P1D8) or a rabbit pAb (Novus). β-actin is a loading control.

However, most interestingly, many amino acid substitutions caused much higher Rif^R^ mutation frequencies, including predicted loop 1, loop 7, and α6-helix residues (loop 1 R18E and R20E, loop 7 H114A and W115A, and α6-helix R171E, A172E, I173A/E, R175E, R176E, R179E). Moreover, in a few instances, combinations of these amino acid substitutions led to mutation frequencies nearly 100-fold higher than wild-type A3H (*e.g.*, R175E/R176E; quantification of 3 independent experiments in **Figure 3C**). One triple mutant combination even proved toxic to *E. coli* (W115A/R175E/R176E), suggesting it elicits the highest overall DNA mutator activity [toxicity has been reported for wild-type human A3A and A3B (Burns et al., 2013; Stenglein et al., 2010)]. These hypermutator phenotypes support the second model described above in which the RNA binding site in A3H is distinct from the single-stranded DNA binding and catalytic region.

### DNA Deaminase Activity of APOBEC3H and Derivatives in 293T Extracts With and Without RNase A Treatment

To extend the bacterial mutation results, the entire panel of A3H constructs was expressed in human 293T cells and tested for single-stranded DNA cytosine deaminase activity in the absence or presence of RNase A treatment. Human 293T cells have no or very low levels of APOBEC enzymes, and lysates from these cells have no detectable single-stranded DNA cytosine deaminase activity unless transfected with an active APOBEC expression construct [*e.g.*, (Carpenter et al., 2012; Stenglein et al., 2010; Thielen et al., 2007)]. As expected, wild-type A3H and most mutant derivatives show no activity in the absence of RNase treatment (**Figure 4A**, upper panels). However, most of the *E. coli* hypermutators display clear DNA cytosine deaminase activity under these conditions, including loop 1 R18E, loop 7 H114A and W115A, and α6-helix A172E, R175E, and R176E, consistent with an RNA binding defect. Moreover, combinations of activating amino acid substitutions resulted in enzymes with further enhanced DNA deamination activity (*e.g*., R175E/R176E). It is also notable that several of the hypermutators have low steady-state expression levels, suggesting they may even be more active than indicated by this series of experiments (**Figure 4A**, lower panels).

In contrast, nearly all of the constructs showed strong DNA cytosine deaminase activity in the presence of RNase treatment, except those predicted to be defective in binding singlestranded DNA or catalyzing C-to-U deamination, which also complements the bacterial mutation data (*e.g.*, catalytic E56A, loop 1 R21E, P22A, Y24A, and P25A, and loop 7 R110E, L111A, Y112A, and Y113A; **Figure 4B**). Immunoblots of the same cell extracts also showed that most constructs, apart from several hyperactive mutants (as above), are expressed at near wild-type A3H levels. It is further notable that these expression controls were done with two different anti-A3H antibodies to avoid having to use potentially confounding heterologous epitope tags and to mitigate the possibility of amino acid changes causing alterations in antibody binding. For instance, A3H constructs with changes spanning residues Lys16 to Arg20 were detected readily using a rabbit polyclonal reagent, but showed no signal with our mouse anti-A3H monoclonal antibody P1D8, consistent with prior data mapping the binding site of this antibody to the N-terminal 30 residues (Refsland et al., 2014). Overall, the results of the Rif^R^ mutation experiments and *in vitro* DNA deaminase studies correlate strongly, with several hyperactive mutants standing-out in both assays.

### Human APOBEC3H Purification and X-ray Structure Determination

To directly compare the RNA binding activities of wild-type A3H and a representative hyperactive mutant, we expressed His6-mCherry-tagged A3H constructs in *E. coli* and performed metal affinity purification. Again, wild-type A3H required RNase A to bind to the affinity resin, whereas the hyperactive mutant did not (**Figure 5A** and similar data not shown for additional hyperactive mutants). Wild-type A3H was further purified using ion exchange column followed by size exclusion chromatography, and it eluted at a size consistent with dimerization (~100 kDa) (**Figure 5B**). This wild-type preparation also had a high 260/280 ratio (>1) suggesting some RNA may still be bound. In contrast, the hyperactive mutant eluted as a monomer with a low 260/280 ratio (~0.6) consistent with near-pure protein (**Figure 5B**). These biochemical results support a model in which RNA mediates dimerization of wild-type A3H.

**Figure 5.**
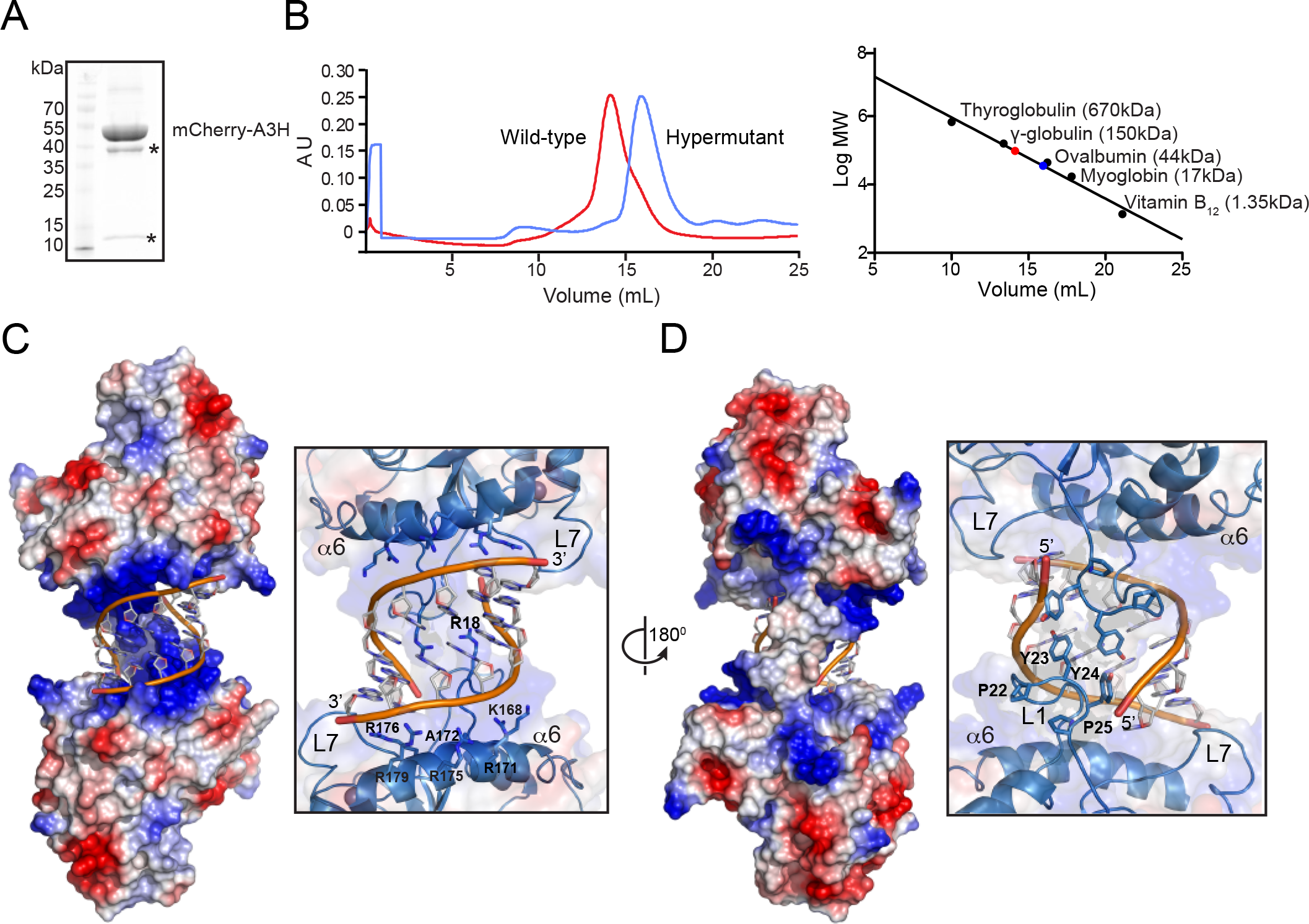
Structure of a human APOBEC3H Dimer Bound to a RNA Double Helix. (**A**) Coomassie-stained gel image showing mCherry-A3H-K52E recovery from *E. coli*. The asterisks represent known heat-induced mCherry degradation products (Gross et al., 2000). (**B**) SEC profiles of wild-type A3H and a representative RNA binding-defective mutant (W115A/R175E/R176E with E56A to prevent toxicity in *E. coli*). Molecular weight standards are shown to the right. (**C**) X-ray structure of human A3H in complex with duplex RNA. Left: a crystallographic pose showing the A3H electrostatic surface potential and double-stranded RNA sandwiched between the positively charged surfaces of two A3H monomers (*i.e*., patch 2). Center: a partially transparent rendering of the same pose highlighting loop 7 (L7) and α6-helix. Right: a zoom-in of the duplex RNA-binding region highlighting positively charged residues in loop 1 (L1 R18) and α6-helix (K168, R171, A172, R175, R176, and R179). (**D**) A 180 degree rotation of the crystallographic poses in panel C to show the back side of the positively charged cradle in which double-stranded RNA is bound (loop 7, L7, is labeled for orientation). The zoom-in depicts the observed loop 1 contacts between the two A3H monomers. See text for additional details.

We next sought to determine the structure of wild-type human A3H to provide a molecular explanation for the aforementioned genetic and biochemical data. During purification from *E. coli* we found that wild-type A3H is prone to aggregation and precipitation. We therefore tried multiple N-terminal solubility tags (Sumo, MBP, GST, GFP, mCherry) and found that mCherry-A3H is soluble but fails to crystallize. We then optimized the linker region between mCherry and A3H, and this resulted in non-diffracting crystals. As discussed above, the A3H surface is predicted to be positively charged with several Lys and Arg residues projecting into solution, and therefore we reasoned that an excess concentration of positive charge may hinder crystallization. In an effort to preserve patch 2 (**Figure 2A**), we examined changes in loop 3 Lys residues (Lys50-Lys51-Lys52). A K52E variant shows DNA editing activity similar to wild-type A3H by both Rif^R^ and *in vitro* activity assays (above), indicating fully intact DNA or RNA binding activities. We therefore changed Lys52 to Glu and used the resulting active enzyme – mCherry-A3H-K52E – for purification, crystallization, and structure determination.

The structure revealed two A3H monomers bridged by a RNA double helix (**Figures 5C** and **5D**; **Supplementary Figure S1**). Each A3H monomer has a cytidine deaminase fold with a central 5-strand β-sheet and 6 surrounding α-helices (**Supplementary Figure S2**). A single zinc ion is coordinated in the active site of each monomer. The 7 base pair RNA duplex with an additional 1 nucleotide overhang is nestled between the same region of each A3H monomer, which is anchored by positively charged α6-helix residues: Arg171, Arg175, Arg176, and Arg179. These residues make direct interactions with the RNA phosphate backbone and encompass a total surface area of 650 Å^2^ (**Figure 5C**). Notably, Arg to Glu substitutions at all of these positions caused increased activity in both the Rif^R^ mutation assay and the *in vitro* DNA deamination assay, with combinations eliciting the highest activity (*e.g.*, R175E/R176E; **Figures 3** and **4**). In addition, although Ala172 does not appear to interact directly with the RNA, an A172E substitution yields a hyperactive protein (RNA binding-defective), likely due to steric and repulsive electrostatic effects.

Additional A3H-RNA contacts are made by loop 1 and 7 residues, as anticipated by the *E. coli* and *in vitro* activity data. In particular, Trp115 of loop 7 is base stacked with the 5’ end of the RNA, and a portion of loop 1, including Arg18, of each monomer projects into the major grove of the RNA helix (**Figure 5C**). The Tyr residues in the loop 1 PYYP motif make direct contacts with the RNA phosphate backbone, and a Tyr23 interaction may also facilitate a monomer-monomer contact (**Figure 5D**). Overall, loop 1, loop 7 and α6-helix residues cradle the RNA duplex through extensive interactions. These data demonstrate a mechanism in which duplex RNA is the prime mediator of A3H dimerization.

### RNA-binding Is Required for APOBEC3H Cytoplasmic Localization and HIV-1 Restriction Activities

As reviewed in the **Introduction**, an APOBEC-RNA interaction is thought to have a role in subcellular localization and to be the major mechanism responsible for the preferential packaging of these deaminases into HIV-1 particles, which is an essential step in the overall retrovirus restriction mechanism (Harris and Dudley, 2015; Malim and Bieniasz, 2012; Simon et al., 2015). Previous studies have shown that A3H is predominantly cytoplasmic, and that this activity is conserved between the human and rhesus macaque enzymes (Hultquist et al., 2011; Li and Emerman, 2011). To test whether the RNA binding domain of A3H is required for cytoplasmic localization, we performed a series of fluorescent microscopy experiments in 293T cells with untagged, wild-type human A3H and a panel of RNA binding mutants. As controls we used cytoplasmic A3G and nuclear A3B. As expected, wild-type A3H is predominantly cytoplasmic. In contrast, all RNA binding domain mutants distribute cell-wide and fail to concentrate in the cytosol (representative images in **Figure 6A** and quantification in **Figure 6B**). Taken together with the aforementioned genetic, biochemical, and structural data, these results strongly indicate that RNA binding is an essential part of the A3H cytoplasmic localization mechanism.

**Figure 6.**
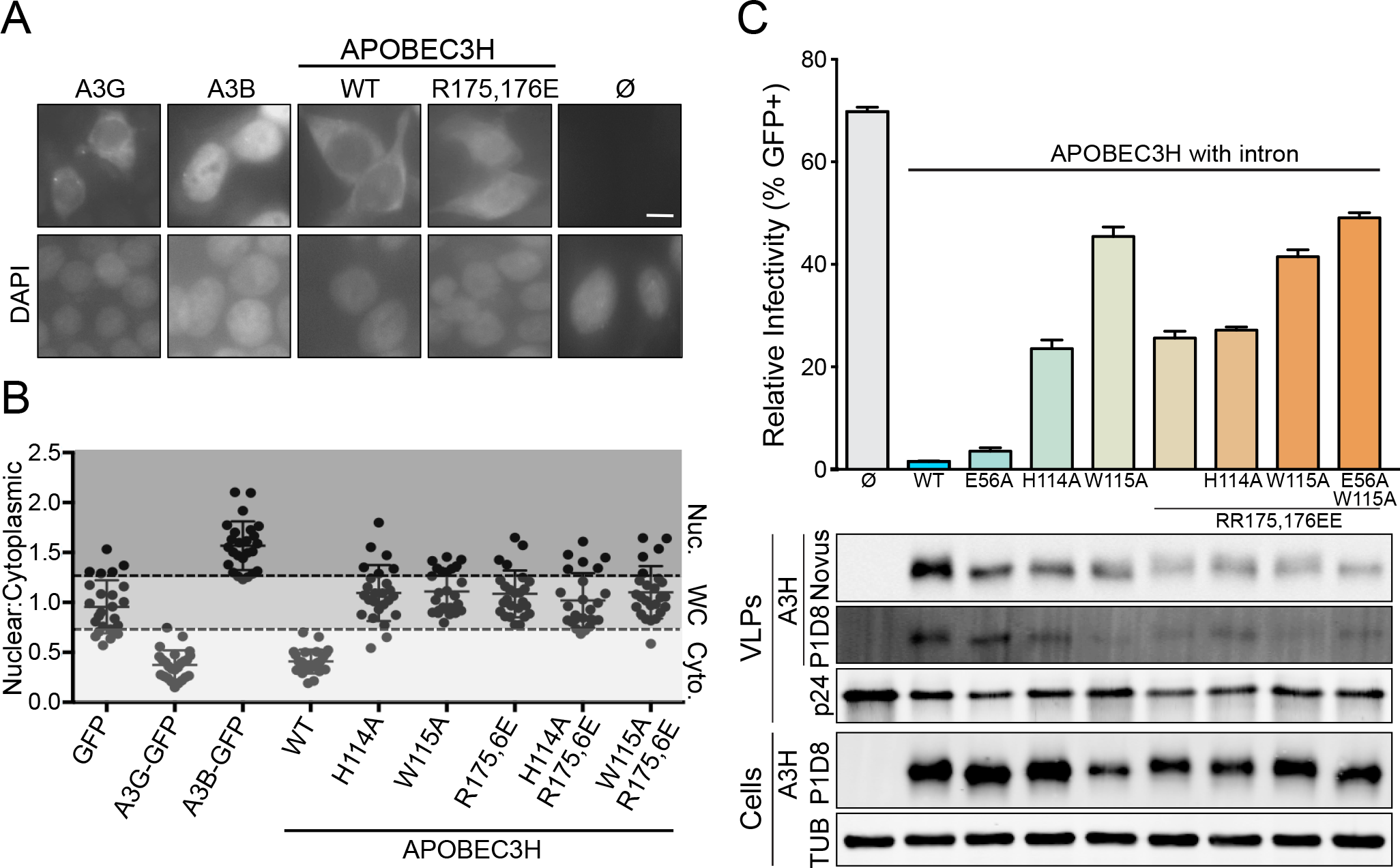
RNA-binding Is Required for APOBEC3H Localization and HIV-1 Restriction. (**A**) Representative images of A3H, A3H-R175E/R176E, A3B, and A3G in 293T cells. The no APOBEC control shows low background fluorescence. DAPI staining defines the nuclear compartment (10 μm scale bar). (**B**) Quantification of the subcellular localization of the indicated constructs (n =25 cells with average +/− SD; N, nuclear; WC, whole cell; C, cytoplasmic). (**C**) Single-cycle infectivity data for Vif-null HIV-1_LAI_ produced in the presence of 200 ng of the indicated untagged, human A3H expression constructs (WT, wild-type; average +/− SD of 3 technical replicates). Corresponding immunoblots are shown below for proteins in virus-like particles (VLPs) and virus-producing 293T cells (A3H detection with mouse mAb P1D8 and rabbit pAb Novus; anti-p24 and anti-tubulin are loading controls for VLPs and cells, respectively).

We next asked whether the RNA binding activity of A3H is required for HIV-1 encapsidation and restriction. Single-cycle infectivity experiments were performed using 293T cells to produce Vif-deficient HIV-1 in the presence of wild-type A3H, a catalytic mutant E56A, or the same RNA binding-defective constructs used above in localization studies (**Figure 6C**). As expected, wild-type A3H causes a >10 fold decline in viral infectivity compared to no APOBEC (vector only) control, and the catalytic mutant E56A exerts a similar suppressive effect [*e.g.*, (Mitra et al., 2015)]. This suggests that most of the anti-HIV-1 activity of A3H is independent of DNA deaminase activity. In contrast, all of the RNA binding-deficient A3H mutants elicited much less than wild-type antiviral activity. For instance, in comparison to the no APOBEC (vector only) control, the R175E/R176E double mutant only caused a 2-fold reduction in antiviral activity. This effect could be further weakened with an additional RNA binding domain substitution (H114A or W115A), and slightly more by adding E56A, further substantiating a large role for RNA binding activity versus a comparatively minor role for DNA deaminase activity. The corresponding immunoblots indicate that most of the compromised antiviral activity of the RNA binding mutants is due to defective packaging into viral particles. Collectively, our results show that the RNA binding activity of A3H is critical for both cytoplasmic localization and HIV-1 restriction mechanisms.

## DISCUSSION

Human A3H is unique among the APOBEC3 proteins with HIV-1 restriction activity because it is the only enzyme with a single zinc-coordinating domain (A3D, A3F, and A3G have double-domain architectures). Thus, it has to pack more innate immune and biochemical activities into a smaller protein unit. Here, we show that A3H binds RNA through a novel RNA duplex-mediated dimerization mechanism. The RNA binding domain was initially identified through mutagenesis experiments, which yielded a group of A3H mutants with *E. coli* hypermutator activity, RNase-independent DNA deaminase activity in 293T cell extracts, and monomeric size exclusion profiles. The first x-ray crystal structure of the human A3H enzyme demonstrates that all of the amino acid substitutions that confer these unique properties map to the protein surface involved in binding to RNA. The physiological importance of the A3H RNA binding domain is evidenced by subcellular localization and HIV-1 restriction experiments in which RNA binding-defective mutants are compromised for cytoplasmic localization, packaging into viral particles and restricting virus infectivity (despite being hyperactive DNA deaminases). An additional surprise from these HIV-1 restriction experiments is the observation that A3H RNA binding activity is more important than DNA deaminase activity. Our results therefore provide molecular and structural explanations for the overall model for HIV-1 restriction in which an RNA interaction mediates A3H cytoplasmic localization, packaging into viral particles and, post-encapsidation, a block to HIV-1 reverse transcription (likely by binding to structured viral genomic RNA elements).

The majority of the restriction activity for the related HIV-1 restriction factor A3G is deaminase-dependent (Browne et al., 2009; Miyagi et al., 2007; Schumacher et al., 2008). In contrast, the majority of A3H antiviral activity is deaminase-independent and, based on results here, most likely due to a direct RNA-binding mechanism. This conclusion is consistent with data showing that wild-type A3H and catalytic mutant E56A similarly suppress HIV-1 reverse transcription *in vitro* (Mitra et al., 2015). Therefore, an important future question is what subset of cellular structured (duplex) RNA species is normally bound by A3H. For example, the abundant Alu-like 7SL RNA has been shown to interact with A3H and may contribute to packaging into HIV-1 particles (Zhen et al., 2012). Moreover, RNA CLIP-seq experiments indicate that A3H binds cellular RNA species relatively non-specifically but adopts preferential partners within mature virions (*e.g.*, ~70% structured regions within HIV-1 genomic RNA and ~20% 7SL RNA) (York et al., 2016). Considering that catalytically dead A3H-E56A still strongly restricts HIV-1, it is likely that HIV-1 genomic RNA is a *bona fide* biological target of A3H. Moreover, the extensive contacts of the α6-helix and loop 1 along the RNA backbone and major groove of the RNA double helix support a mechanism in which A3H recognizes specific RNA structures. The precise nucleobase sequences of these RNA structures will also be of future interest because, despite clear A-form RNA in our A3H x-ray structure (**Supplementary Figure S1**), the crystal lattice has a mixed population of RNA species derived from *E. coli*.

Amino acid alignments strongly indicate that the human A3H-RNA binding mechanism is conserved among primate A3H enzymes, and likely also for homologous mammalian Z3-type deaminases (**Supplementary Figure S3**). In particular, residues homologous to human A3H loop 1 Arg 18, loop 7 His114 and Tyr115, and α-helix 6 Arg171, Ala172, Arg175, Arg176, and Arg179 are present in all primate A3H enzymes. Mammalian A3Z3 homologs also have near-identical α-helix 6 and loop 7 residues, as well as positively charged residues situated toward the N-terminal end of loop 1. More distantly related APOBEC family members may also have similar duplex RNA binding mechanisms. For instance, human A3C (Z2-type enzyme) and human AID have analogous Arg-rich α6 helices and loop 1 motifs. Interestingly, AID was shown recently to bind G-quadraplex DNA structures using this basic region (Qiao et al., 2017). Although duplex RNA was not tested in these studies, given the demonstrated biochemical association between AID and RNA (Bransteitter et al., 2003), it is conceivable that the same region may bind structured RNA, perhaps as part of an R-loop directed DNA targeting mechanism relevant to antibody diversification. Indeed, a variety of AID α6 mutants are defective in class switch recombination, including naturally occurring hyper-IgM disease alleles (Durandy et al., 2007). Other single domain enzymes such as APOBEC1 and A3A lack similar RNA binding residues and therefore may bind both RNA and single-stranded DNA using active site residues and surrounding loop regions (especially loops 1 and 7). This inference is supported by the fact that both of these enzymes are RNA editing enzymes, potent DNA deaminases, and, at least for A3A, largely monomeric in single molecule studies *in vitro* (Shlyakhtenko et al., 2014) and in living human cells where RNA is abundant (Li et al., 2014). The double domain enzymes, A3B, A3D, A3F, and A3G, are also subject to regulation by RNA (**Introduction**) and further work will be needed to assess conservation with the mechanism described here.

The roles of RNA in regulating the multifaceted functions of the APOBEC/AID family of polynucleotide editing enzymes have been difficult to define mechanistically, in large part due to a lack of separation-of-function mutants and structural information. The studies presented here with human A3H demonstrate a novel duplex RNA binding mechanism through identification and analyses of separation-of-function mutants and the determination of an A3H-duplex RNA co-crystal structure. The negative regulatory role of RNA in suppressing the A3H DNA deaminase activity, as well as the positive regulatory role of RNA in governing cytoplasmic localization, may both be relevant to preventing the accumulation of genomic mutations. As A3H is a likely contributor to the overall APOBEC mutation signature in cancer (Starrett et al., 2016), it will be interesting to ask whether somatic mutations in A3H or other cellular factors (RNA or protein) destabilize its RNA binding activity and thereby contribute to tumor evolution. Most importantly, here, the RNA binding activity of A3H is essential for cytoplasmic localization and HIV-1 restriction. Given the fact that 5 different A3H hypermutators are defective for RNA binding and cytoplasmic localization, it is likely that the same duplex RNA binding mechanism is responsible for both localization and packaging into HIV-1 virions. Moreover, once encapsidated, the strong RNA binding activity of A3H likely blocks reverse transcription, as demonstrated by analyses of the E56A catalytic mutant. Thus, the mechanistic and structural studies here shed light on this deaminase-independent HIV-1 restriction mechanism and strongly suggest that this strong RNA binding activity will prove relevant to suppressing the spread of multiple other viruses and transposons.

## Methods

### Plasmids for APOBEC3H Expression in *E. coli*

6xHis-SUMO-A3H was generated by PCR amplifying A3H HapII 1-183 cDNA (GenBank: ACK77775.1) from pcDNA3.1-A3H (Hultquist et al., 2011) using primers 5’-gcgcGGTCTCTAGGTGGCGGCGGCATGGCTCTGTTAACAGCCG and 5’-gcgcTCTAGATTAGGACTGCTTTATCCTC. The PCR product was digested with BsaI and XbaI and ligated into similarly digested pE-SUMO (Life Sensors). This forward primer also encodes a triple Gly linker.

His-tagged mCherry(1-232)-3xAla linker-HapII(1-183) was generated by overlap PCR using primers 5’-GGATCCGAATTCGCTGGAAGTTCTGTTCCAGGGGATGGTGAGCAAGGGCGAGGAGG, 5’-CTCCACCGGCGGCATGGACGCCGCTGCCATGGCTCTGTTAACAGCCGAAACATTCC, 5’-GGAATGTTTCGGCTGTTAACAGAGCCATGGCAGCGGCGTCCATGCCGCCGGTGGAG, and 5’-GCGCGCGGCCGCTCAGGACTGCTTTATCC. The PCR product was digested with EcoRI and NotI and ligated into MCS1 of the prSFDuet-1 (Novagen). A HRV rhinovirus 3C protease site was engineered prior to the mCherry ATG start site (sequence included in the forward primer used during overlapping PCR). A3H-K52E was introduced by site-directed mutagenesis. The mCherry coding sequence was amplified from pRSETB-mCherry (Shaner et al., 2004) and A3H was amplified from His-SUMO-A3H (above). Single amino acid substitution derivatives were generated using site-directed mutagenesis, and Sanger sequencing was used to verify all constructs.

### *E. coli*-based Rif^R^ Mutation Assays

Experiments were done as described (Harris et al., 2002; Shi et al., 2015). His6-SUMO-A3H constructs were transformed into *E. coli* C43(DE3) cells, and single colonies were grown to saturation at 37°C in 2 mL LB plus ampicillin (100 μg/ml). A 100 μl aliquot of undiluted culture was plated directly onto LB-agar plates containing rifampicin (100 μg/ml) to select Rif^R^ mutants. Cultures were diluted serially in 1X M9 salts (sodium phosphate dibasic, potassium phosphate dibasic, sodium chloride, ammonium chloride) and plated on LB to determine viable cell counts. Mutation frequencies were calculated by dividing the number of colonies on the rifampicin plate by the total number of cells in the overnight culture (total number colonies on LB plates and multiplied by the dilution factor).

### Plasmids for A3H Expression in Human Cell Lines

Coding sequences of A3H exon 2 and exons 3-5 were amplified from pcDNA3.1-A3H (Hultquist et al., 2011) using primer pairs 5’-NNNNGGTACCACCATGGCTCTGTTAACAG/5’-AAACATCTCCTGGACTCACCTTGTTTTCAAAGTAGCCTC and 5’-GTCTCCTTTCATCTCAACAGAAAAAGTGCCATGCGGAAAT/NNNNGCGGCCGCTCAGGACTGCTTTATCCT, respectively. Human *HGB2* intron 2 was amplified from an A3Bi containing plasmid (Burns et al., 2013) using primer pair 5’-GAGGCTACTTTGAAAACAAGGTGAGTCCAGGAGATGTTT/5’-ATTTCCGCATGGCACTTTTTCTGTTGAGATGAAAGGAGAC. The amplified fragments were fused together by overlap extension PCR, and inserted into the KpnI-NotI cloning site of pcDNA3.1(+) (ThermoFisher Scientific). Mutant derivatives of pcDNA3.1-A3Hi were generated using site-directed mutagenesis (primers available on request), and Sanger sequencing was used to verify the integrity of all plasmids.

### APOBEC3H DNA Deamination Experiments

293T cells were plated in 6-well plates at 400,000 cell density. The following day, the cells were transfected with 1 μg of each pcDNA3.1-A3Hi expression construct and 50 ng of an eGFP expression plasmid (transfection control). 48 hrs later, cells were harvested and resuspended in 200 μL HED buffer (25 mM HEPES pH 7.4, 15 mM EDTA, 1 mM DTT, Roche Complete EDTA-free protease inhibitors, 10% glycerol) and frozen at -80°C. Cells were thawed, vortexed, sonicated for 20 min in a water bath sonicator (Branson), and centrifuged 10 min at 16,000 g, and then soluble extracts were transferred to a new tube. Half of each extract was treated with RNase A (100 μg/ml) at RT for 1 hr. An oligo master mix containing 1.6 μM 5’-ATTATTATTATTCTAATGGATTTATTTATTTATTTATTTATTT-fluorescein in HED buffer was made and mixed 1:1 with cell extracts (10 μL). Reactions were gently mixed, spun down, then incubated at 37°C for 60 minutes, heated to 95°C for 10 minutes to stop the reactions, then incubated with 0.1 U/rxn UDG for 10 minutes at 37°C. Sodium hydroxide was added to a final concentration of 100 mM and reactions were heated to 95°C to cleave the DNA at abasic sites. Reactions were mixed with 2x DNA PAGE loading dye (80% formamide, 1x TBE, bromophenol blue, xylene cyanol). Reaction products were separated by 15% denaturing PAGE (or TBE-Urea PAGE) and visualized by scanning on a Typhoon FLA-7000 scanner on fluorescence mode.

A fraction of each extract was separated by 12% SDS-PAGE and transferred to low-fluorescence PVDF overnight at 20 V. Membranes were blocked with 4% milk in PBST then incubated with mouse anti-A3H mAb P1D8 (Refsland et al., 2014) or rabbit anti-β-actin pAb (Cell Signaling: 13E5), at dilutions of 1:1,000 or 1:10,000 respectively. After washing in PBST, membranes were probed with the goat anti-mouse 680 antibody (Life Technologies: A21057) or the goat anti-rabbit 800CW (Li-cor Biosciences: 926-32211), each 1:20,000. Blots were stripped and reprobed with rabbit anti-A3H pAb at 1:1,000 (Novus Biologicals NBP1-91682), followed by Li-cor anti-rabbit 800CW secondary at 1:20,000. Imaging was done with a Li-cor Odyssey.

### APOBEC3H purification from *E. coli*

His6-mCherry-A3H HapII 1-183 was transformed into OneShot Bl21(DE3) competent cells (Thermo-Fischer Scientific) and, after overnight incubation, colonies were directly inoculated into 2XYT media containing 50 μg/ml kanamycin. Approximately 30min. prior to induction, cultures were supplemented with 100 μM zinc sulfate and cooled to 16°C. Protein expression was induced overnight by addition of 0.5 mM IPTG at ~1 OD_600_. Cells were centrifuged 3800 g for 20 min and resuspended in lysis buffer containing 50 mM Tris pH 8, 500 mM NaCl, 5 mM imidazole, lysozyme, and RNase A (10 mg). Cells were then sonicated (Branson Sonifer) and cell debris was removed by centrifugation (13000g, 45 min). The supernatant was added to Talon Cobalt Resin (Clontech), washed extensively with wash buffer (50 mM Tris-HCl pH 8, 500 mM NaCl, 5mM imidazole), and eluted with elution buffer (50mM Tris-HCl pH 8, 500 mM NaCl, 250 mM imidazole). Fractions containing protein were pooled and dialyzed overnight into buffer containing 50 mM Tris-HCl pH 8, 100 mM NaCl, 5% glycerol, 2mM DTT. The protein was concentrated to ~5mLs and loaded onto a 5mL HiTrap MonoQ cartridge (GE Healthcare) equilibrated with wash buffer (50 mM Tris-HCl pH 8, 100 mM NaCl). The bound protein was washed with wash buffer and eluted with a linear gradient of high salt buffer (50 mM Tris pH 8, 1 M NaCl). The protein elutes between 30-40% high salt buffer. Fractions were collected and purified further with a 26/600 S200 Gel Filtration column (GE Healthcare) using a buffer contain 20 mM Tris-HCl pH 8 and 500 mM NaCl. Fractions containing purified protein were pooled, mixed with 5 mM DTT, and concentrated to 25 mg/ml. The final purified protein had an OD280/260 measurement greater than 1, indicating bound nucleic acid.

### APOBEC3H Crystallization

Prior to crystallization, 10 mM DTT and trace amounts of human rhinovirus 3C protease was added to an aliquot of protein diluted to 20 mg/ml to remove the 6xHis tag. A3H was crystallized using the sitting drop vapor diffusion in a solution containing 18% PEG 3350 (v/v) and 300 mM ammonium iodide. The crystal trays were incubated at 18°C and protein crystals appeared overnight and grew to full size within 2-5 days. The crystals were preserved in cryprotectant solution containing 20% PEG3350, 300 mM ammonium iodide, and 20% glycerol.

### X-ray Data Collection and Structural Determination

Initial x-ray diffraction data were collected at the Advanced Photon Source NE-CAT 24-ID-E beamline at the wavelength of 0.979 Å. The datasets were processed using HKL2000 (Otwinowski and Minor, 1997) and XDS (Kabsch, 2010), initially in the space group P6_1_22.Structure solution by molecular replacement was unsuccessful despite exhaustive attempts using mCherry and various A3 structures as search models. Subsequently, high-multiplicity x-ray diffraction data were collected at the NE-CAT 24-ID-C beamline at the zinc K-edge wavelength of 1.282 Å. Single-wavelength anomalous dispersion (SAD) phasing using SHELX C/D/E (Sheldrick, 2010) and PHENIX (Adams et al., 2010) produced an electron density map that clearly showed secondary structure elements of A3H and a segment of RNA double-helix, which has a deep and narrow major groove and a shallow and wide minor groove (**Supplementary Figure S1**). The A3H and RNA molecules were built into the experimental map using COOT (Emsley et al., 2010). The *mFo-DFc* difference map after initial refinement with phase-restraints against the SAD-derived phases revealed a β-barrel structure corresponding to partial mCherry. Including mCherry in the refinement lowered the R_free_ by 1.0 %. Refinement in a lower symmetry space group P3_1_12 resulted in significant improvement of R_free_ but refining with a twin operator “-h,-k,l” increased the R_free_. Therefore we continued refinement in the space group P3_1_12 without twinning. Subsequent iterative model building using COOT and refinement with PHENIX suite (Adams et al., 2010) resulted in the final R_work_ / R_free_ of 32.3 / 35.2%. The slightly high R-factors are likely due to the uncertain location of mCherry, which accounts for >50% of the total scattering mass. The model quality during the refinement was constantly checked by combining the SAD-derived and model-based phases to minimize possible model bias. The asymmetric unit contains two A3H molecules, two RNA strands, and two copies of mCherry with occupancy refined to 0.5. However, we modeled two RNA double helices (poly-A paired with poly-U) in the asymmetric unit, each with half-occupancy, to represent the double-stranded RNA molecules comprising mixed population of unknown complementary sequences and following crystallographic 2-fold symmetry as an ensemble. The summary of data collection and model refinement statistics is shown in **Table 1**. Images were produced using PYMOL (http://www.pymol.org). Coordinates have been deposited in the Protein Data Bank with accession code 6B0B.

**Table 1.**
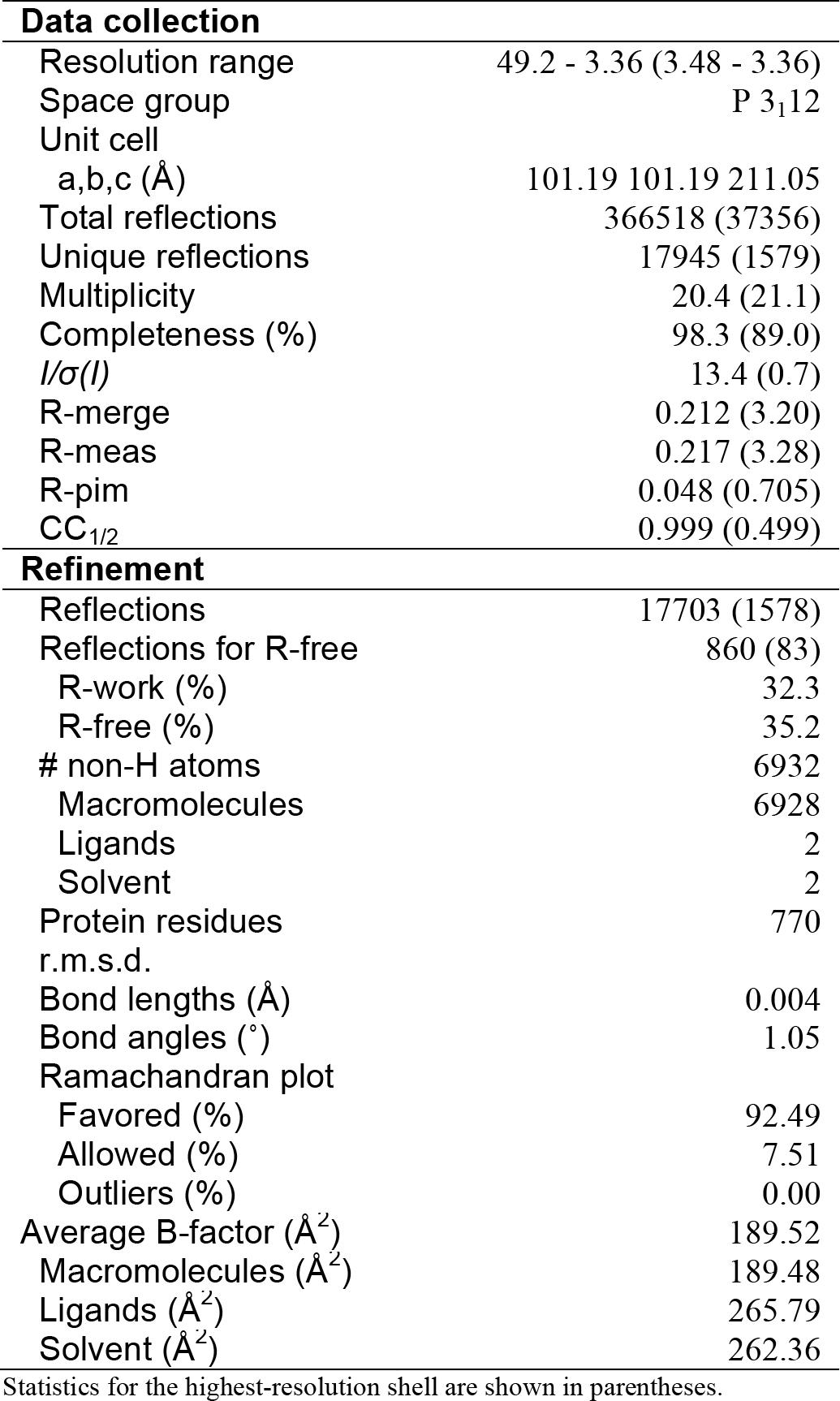
Data collection and refinement statistics.

| **Data collection** |  |
| --- | --- |
| Resolution range | 49.2 - 3.36 (3.48 - 3.36) |
| Space group | P 3<sub>1</sub>12 |
| Unit cell |  |
| a,b,c (Å) | 101.19 101.19 211.05 |
| Total reflections | 366518 (37356) |
| Unique reflections | 17945 (1579) |
| Multiplicity | 20.4 (21.1) |
| Completeness (%) | 98.3 (89.0) |
| *I*/$\sigma$(*I*) | 13.4 (0.7) |
| R-merge | 0.212 (3.20) |
| R-meas | 0.217 (3.28) |
| R-pim | 0.048 (0.705) |
| CC<sub>1/2</sub> | 0.999 (0.499) |
| **Refinement** |  |
| Reflections | 17703 (1578) |
| Reflections for R-free | 860 (83) |
| R-work (%) | 32.3 |
| R-free (%) | 35.2 |
| # non-H atoms | 6932 |
| Macromolecules | 6928 |
| Ligands | 2 |
| Solvent | 2 |
| Protein residues | 770 |
| r.m.s.d. |  |
| Bond lengths (Å) | 0.004 |
| Bond angles (°) | 1.05 |
| Ramachandran plot |  |
| Favored (%) | 92.49 |
| Allowed (%) | 7.51 |
| Outliers (%) | 0.00 |
| Average B-factor (Å<sup>2</sup>) | 189.52 |
| Macromolecules (Å<sup>2</sup>) | 189.48 |
| Ligands (Å<sup>2</sup>) | 265.79 |
| Solvent (Å<sup>2</sup>) | 262.36 |

Statistics for the highest-resolution shell are shown in parentheses.

### Fluorescent Microscopy Experiments

Approximately 10,000 293T cells were plated into a 96-well CellBIND microplate (Corning) and allowed to adhere overnight. Cells were transfected with 100 ng of the indicated untagged A3H construct. For localization controls, 100 ng of A3B-eGFP, A3G-eGFP, or eGFP expression vector were transfected. 48 hr post-transfection, cells were washed twice with PBS and fixed with 4% paraformaldehyde (20 min, RT). Cells were then washed twice with cold PBS and permeabilized with 0.5% Triton-X100 (10 min, 4°C). Cells were washed once with cold PBS, incubated in blocking buffer (4% BSA, 5% goat serum, in PBS) for 1 hr at RT, and then stained with 1:100 rabbit anti-A3H pAb (Novus Biologicals, NBP1-91682), or with a rabbit anti-A3B mAb (5210-87-13) at 1:500 that also cross-reacts with A3G (Leonard et al., 2015). Cells were washed with cold PBS and stained with 1:1000 goat anti-rabbit Alexaflour 594 (Invitrogen, A11037) for 1 hr at RT. After additional washing, cells were incubated in PBS containing 0.1% DAPI. Images were collected at 40x magnification using an EVOS FL Color microscope (ThermoFisher),and quantified using ImageJ software. Individual cells were scored and grouped into three categories (nuclear, whole cell, or cytoplasmic). Subcellular compartment were defined based on DAPI staining (nucleus). The anti-A3 signal was calculated by dividing the nuclear anti-A3 staining intensity by the cytoplasmic intensity, and then graphed with Prism 6.0 (GraphPad Software).

### HIV-1 Restriction Experiments

Tandem stop codons were introduced into codons 26 and 27 of the full-length HIV-1 molecular clone pLAI.2 (Peden et al., 1991) (AIDS Reagent Program 2532) by subcloning the *vif-vpr* region into pJET1.2, performing site-directed mutagenesis (Stratagene), and shuttling the *vif-vpr* region back into the original vector using PshAI and SalI restriction sites. 50% confluent 293T cells were transfected (TransIt, Mirus) with 1 μg Vif-deficient pLAI.2 and 200 ng pcDNA3.1-A3Hi (above) or 200 ng empty vector. After 48 hr incubation, viral supernatants were cleared of cells by filtration (0.45 μm) and used to infect CEM-GFP cells to monitor infectivity via flow cytometry.

Cell and viral particle lysates were prepared for immunoblotting as follows. Cells were pelleted, washed, and then lysed with 2.5X Laemmli sample buffer. Virus containing supernatants were filtered and virus-containing supernatants were pelleted via centrifugation through a 20% sucrose cushion, then lysed in 2.5X Laemmli sample buffer. Lysates were then subjected to SDS-PAGE followed by protein transfer to PVDF using a Bio-Rad Criterion system. Membranes were probed with a mouse anti-A3H mAb P1D8 (Refsland et al., 2014), a rabbit anti-A3H pAb (NBP1-91682, Novus), an anti-HIV-1 p24/CA mAb (AIDS Reagent Program 3537), and a mouse anti-tubulin mAb (Covance). Secondary antibodies were goat anti-rabbit IRdye 800CW (Li-cor 926-32211) and goat anti-mouse Alexa Fluor 680 (Molecular Probes A21057). Membranes were imaged using a Li-cor Odyssey instrument, and images were prepared for presentation using Image J 1.49 (http://rsb.info.nih.gov/ij/download.html).

## ACKNOWLEDGMENTS

We thank Rena Levin-Klein for providing the wild-type A3H construct and JingYing Zhang for technical assistance. The following reagents were obtained through the NIH AIDS Reagent Program, Division of AIDS, NIAID, NIH: pLAI.2 from K. Peden, courtesy of the MRC AIDS Directed Program, and anti-HIV-1 p24/CA mAb courtesy of B. Chesebro and K. Wehrly. This work was supported by grants from the US National Institutes of Health (NIGMS R01-GM118000 to RSH and HA, NIGMS R35-GM118047 to HA, NCI R21-CA206309 to RSH), the Prospect Creek Foundation (RSH), and the University of Minnesota College of Biological Sciences, Academic Health Center, and Masonic Cancer Center. This work is based upon research conducted at the Northeastern Collaborative Access Team beamlines, which are funded by the US National Institutes of Health (NIGMS P41-GM103403). The Pilatus 6M detector on 24-ID-C beamline is funded by a NIH-ORIP HEI grant (S10 RR029205). This research used resources of the Advanced Photon Source, a U.S. Department of Energy (DOE) Office of Science User Facility operated for the DOE Office of Science by Argonne National Laboratory under Contract No. DE-AC02-06CH11357. RSH is the Margaret Harvey Schering Land Grant Chair for Cancer Research, a Distinguished University McKnight Professor, and an Investigator of the Howard Hughes Medical Institute.

### AUTHOR CONTRIBUTIONS

NMS and RSH conceived and designed the studies. KVL, MWL, and NMS created all the constructs. KVL and MWL performed bacterial mutation experiments. MAC performed DNA deamination activity assays. NMS purified and crystallized A3H. SB and KS collected and phased diffraction data. KS determined the structure, and KS, NMS, HA analyzed structural data. DJS performed fluorescent microscopy studies. CMR and JW performed HIV-1 restriction experiments. WLB produced antibodies and contributed to project management. NMS and RSH drafted the manuscript, and all authors contributed to figure preparation and revisions.

### COMPETING FINANCIAL INTERESTS

RSH is a co-founder, shareholder, and consultant of ApoGen Biotechnologies Inc. HA is a consultant for ApoGen Biotechnologies Inc. The other authors have no competing financial interests to declare.

